# EXPLICIT CORRECTION OF SEVERELY NON-UNIFORM DISTRIBUTIONS OF CRYO-EM VIEWS

**DOI:** 10.1101/2025.10.07.680880

**Authors:** Charles Barchet, Ottilie von Loeffelholz, Roberto Bahena-Ceron, Bruno P. Klaholz, Alexandre G. Urzhumtsev

## Abstract

The quality of three-dimensional macromolecular image reconstructions by cryo electron microscopy strongly depends on the number and quality of respective two-dimensional projections and on their angular distribution in space. Distributions with one or a few strongly preferred particle orientations may result in maps deformed in certain directions. A simple removal of overrepresented views may improve the quality of the reconstructed maps when the level of noise in the two-dimensional projections is low and the dataset-size can afford this removal but is counterproductive or insufficient otherwise. Giving complementarily an increased weight to underrepresented views, or taking multiple copies of them during the reconstruction, may improve the results, naturally, depending on how much the views distribution is non-uniform. This work describes the results of three-dimensional reconstructions using an explicit correction of the number of over- and underrepresented projections for non-uniformly distributed sets.

**Synopsis:** An explicit numerical leveling of non-uniformly distributed sets of 2D projections with the program VUE improves the 3D reconstructions and illustrate sources of image distortions.

## 1. Introduction

The principal object of a single-particle analysis in cryo electron microscopy (cryo-EM) is a three-dimensional (3D) map of the electrostatic scattering potential, which is obtained by 3D reconstruction from two-dimensional (2D) projections through back-projection (Harauz & van Heel, 1986, as one of the pioneering publications). Such maps are subject of further interpretation, whenever possible, in terms of an atomic model which becomes the norm with the “resolution revolution” (Kühlbrandt, 2014). A cryo-EM map can contain various kinds of errors. Similar to crystallographic analysis (Lunin *et al*., 2002), these errors can be considered either as removable or irremovable. Removable errors are those due to inappropriate choice of the values of the parameters of the data interpretation; these values can be improved. For example, they are errors in the assigned values of the directions under which given 2D projections correspond to the 3D object reconstructed from them. Exact, or at least better values of such orientation parameters, *e*.*g*., of the Euler angles (Heymann *et al*., 2005), can be obtained during refinement. An example of irremovable errors are errors in the experimental 2D projection values themselves recorded during data collection. Programs can treat some of these (*e*.*g*., inactive pixels on the detector, etc.), reduce their influence implementing special protocols, but they cannot be removed.

Missed 2D projections are another example of irremovable errors. It is known for a while that a non-uniform distribution of projections, the views, affects the reconstructed image. Without being exhaustive, we cite Harauz & van Heel (1986), Boisset *et al*. (1998), Unger (2000), Sorzano *et al*. (2021), Baldwin *et al*. (2023), and references therein. The views distributions are analyzed and illustrated with software using different approaches, *e*.*g*., those by Orlov *et al*. (2006), Punjani *et al*. (2017), Grant *et al*. (2018) and a combination of *Relion* (Scheres, 2012) with *Chimera* (Pettersen *et al*., 2004). Measures to characterize the degree of non-uniformity have been proposed (Naydenova & Russo, 2017; Baldwin & Lyumkis, 2021; Urzhumtsev, 2025a). An early approach to account for preferential views has been a correction based on the point-spread function as implemented in the Imagic software during 3D reconstruction (Harauz & van Heel, 1986). Since then other weighting methods (*e*.*g*., Orlov *et al*., 2011; Scheres, 2012; Sorzano *et al*., 2021) have been proposed to correct the respective map distortions.

Recently, we have developed the program *VUE* (Urzhumtseva *et al*., 2024) which transforms such distributions of the views into points on the spherical surface, and projects them in 2D in a quantitatively exact manner using the Lambert projection, thus avoiding spatial distortions, as opposed to other programs. *VUE* allows to detect and visualize view distributions as plots, and to calculate their basic statistics including frequency associated with each individual view, *ν*_*n*_, *n* = 1, …, *N* according to the number of its neighbors and the angular distance to them. Here we present a further development of this program and a novel tool allowing to correct non-uniform distributions of views as much as possible, thus improving the respective 3D reconstructions. This development let us to analyze the effect of such corrections and to search for an optimal protocol. In the following, we explain our approach, describe the tests and their results, giving some examples with simulated and experimental data.

## 2. Method

### 2.1. Overall scheme

Trying to make the distribution of views more uniform means, first of all, to reduce the overrepresented ones as this is done for decades (just as an example, see Shaikh *et al*., 2008, and references therein). Since these views are slightly different and contain different experimental errors, removal of some of them may make some experimental information lost and the 3D reconstruction noisier and more blurred, *i*.*e*., to decrease its resolution.

On the other hand, one needs to complete underrepresented views requiring, formally speaking, an additional experiment. In a certain way, this can be done computationally introducing respective weighting functions during reconstructions as mentioned in the introduction.

Instead, one can simulate this explicitly by repeating several times a reference to a given projection, like if it were recorded several times in exactly the same orientation and containing by chance exactly the same experimental errors. On one hand, this simply repeats the information. On the other hand, this allows to understand better the effects of such overweighting for the underrepresented views. Moreover, one can artificially modify the parameters of a given projection, its view direction, which could give non-zero estimates for some Fourier coefficients missed otherwise. Both these modifications are trivial since they do not change the actual file of projections but only the file containing references to them.

### 2.2. Modification of the set of projections

For most of cryo-EM software, the values of the 2D projections, available at some regular grid, are kept separately from another file containing parameters of these projections, usually a line per projection. This list of parameters includes those describing directions (*e*.*g*., using Euler angles) of the projections with respect to the spatial orientation of the object. The simplest procedure to correct non-uniform distribution consists in removing the overrepresented views. Practically, *VUE* does it by removing respective lines from the list of the projection parameters. These lines are selected randomly, with the probability proportional to the excess over the threshold *ν*_*cut*_, the value chosen by the user. The program *VUE* has an option to keep a slight excess of the overrepresented views, which may contain extra information according to Stagg *et al*. (2014).

A more sophisticated procedure additionally completes the list of projections by extra lines repeating a reference to the given underrepresented projections. This can be seen as the same projection has ‘accidentally’ been measured several times. The number of copies can be taken to increase its frequency up to the same threshold value *ν*_*cut*_, thus equalizing the distribution. The program *VUE* may repeat these references without or with small artificial random perturbation of the view parameters, *i*.*e*., introducing minor angular variances as defined by the user. On the contrary, the projections themselves are not modified.

In this procedure, taking the correction threshold *ν*_*cut*_ too high leaves more overrepresented experimental projections, and this, to equalize the distribution, requires to complete many underrepresented views thus introducing too much ‘dummy’ information affecting the reconstructed image. Inversely, taking *ν*_*cut*_ too low, one reduces the number of the ‘dummy’ projections to be added simultaneously reducing the number of experimental projections for the reconstruction, which may be counterproductive as well.

### 2.3. Experimental data

For our analysis, we used synthetic data, for which the exact answer is known thus simplifying evaluation of the results, and experimental data sets available in the laboratory. We identified three experimental data sets that suffered from a non-uniform views distribution.

First, we considered the cryo-EM data for a human 40S small ribosomal subunit acquired on the onsite G1 Titan Krios cryo electron microscope installed at the CBI. The data set contains about 45,000 2D projections in total. It has a dominating cluster of projections together with a much smaller one at an angular distance about 90 degrees (Fig. 1a).

**Figure 1.**
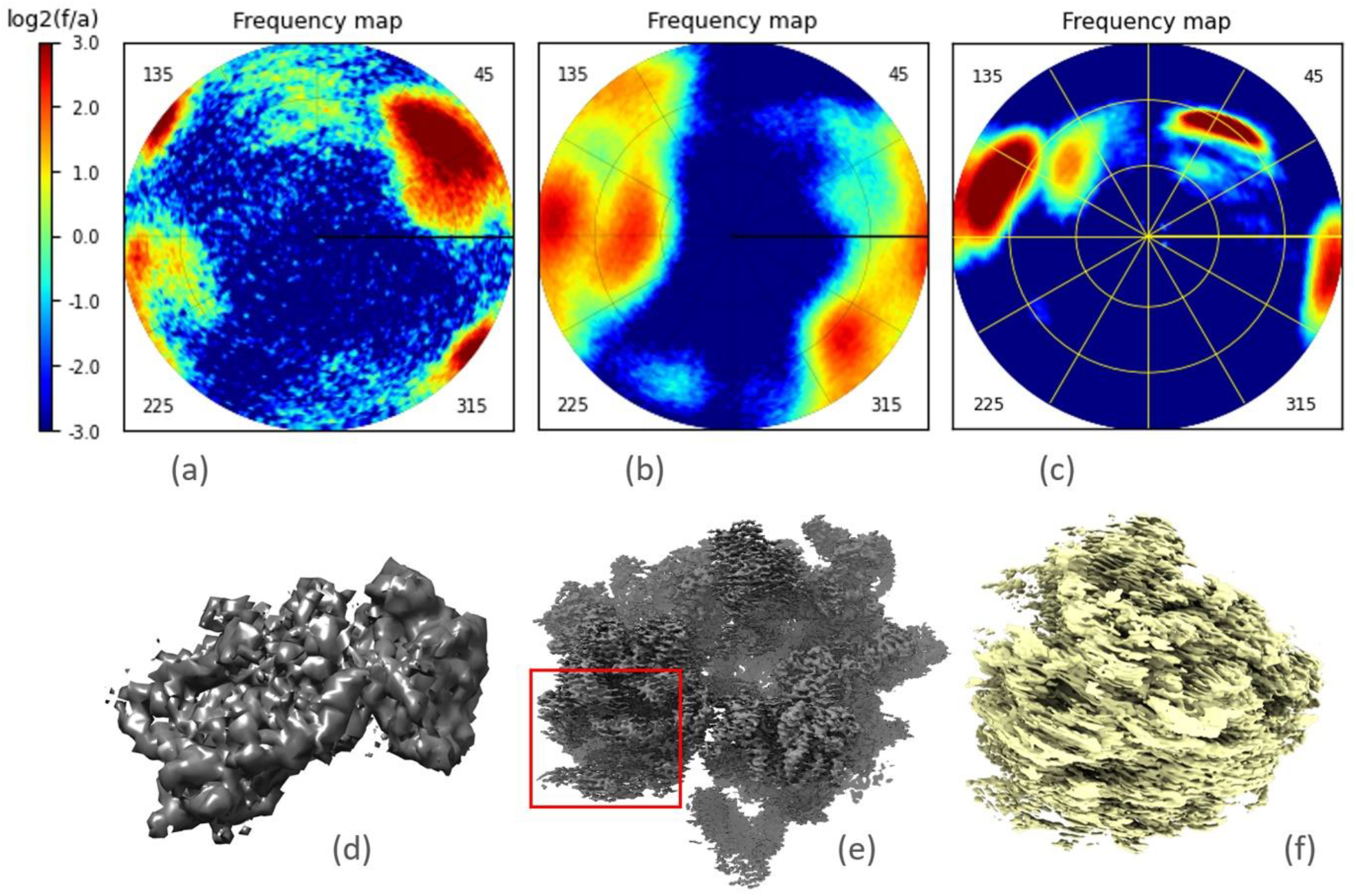
Distribution of views for the experimental data sets used in this work. a) 40S human ribosomal particle; b) 70S *S. aureus* ribosome; c) 80S human ribosome. The color scale is logarithmic (log_2_), with respect to the mean frequency value. d), e) and f) show the maps for the respective 3D reconstructions. The red rectangle in e) indicates the region shown magnified in Fig. 8d. Views distribution and maps were plotted with the programs *VUE* (Urzhumtseva *et al*., 2024) and *Chimera* (Pettersen *et al*., 2004), respectively.

Second, we worked with 2D cryo-EM projections for the 70S *Staphylococcus aureus* ribosome, about 582,000 projections in total. These data have been collected on the in-house G1 Titan Krios cryo electron microscope equipped with a Falcon 4i camera and operating at an acceleration voltage of 300 kV. Figure 1b shows that the views are strongly clustered around three directions relatively distanced by about 45° from each other.

As the third set, we used a cryo-EM data set of the human 80S ribosome the structure of which determined earlier (Khatter *et al*., 2015; Natchiar *et al*., 2017; Holvec *et al*., 2024). The data set were acquired on the in-house G4 Titan Krios cryo electron microscope. The 2D projections, about 337,000 in total, are distributed non-uniformly, with three view clusters, of a different size, concentrated roughly in a plane (Fig. 1c; here left and right blobs are parts of the same cluster).

For each of these data sets, we calculated the 3D reconstruction using the software *Relion* (Scheres, 2012). The map for the small subunit was rather isotropic, with no stripes typical for severely non-uniform distributions (Fig. 1d). The maps for the 70S ribosome showed some deformation (following the horizontal direction in Fig. 1e). The most significant map deterioration was observed for the 80S ribosome (Fig. 1f).

While the first data set was not useful to test modifications of the sets of views, the two latter gave examples for subsequent analysis and expected improvement of maps deteriorated in a different scale.

### 2.4. Simulated data

To quantify the effect of various manipulations with the sets of non-uniformly distributed 2D projections, we have started from a series of tests with simulated data. To facilitate calculations, the relatively small and highly asymmetric structure of initiation factor IF2 was chosen (Simonetti *et al*., 2013). Its atomic model was placed in the center of a P1 cubic unit cell with the edge length equal to 200 Å. The program *Chimera* (Pettersen *et al*., 2004) was used to generate the respective three-dimensional map at a resolution of 2 Å in the grid with a step equal to 1 Å; we refer to it as 2ÅC map. Additionally, we calculated two similar maps at a resolution of 5 Å (5ÅC map) and 10 Å (10ÅC map).

The software *Relion* (Scheres, 2012) was used to generate 100,000 two-dimensional projections of this 3D map, also on the grid with a step size of 1 Å. The directions of these projections were distributed randomly and uniformly in space (Fig. 2a), and we refer to this set as 100Ku.

**Figure 2.**
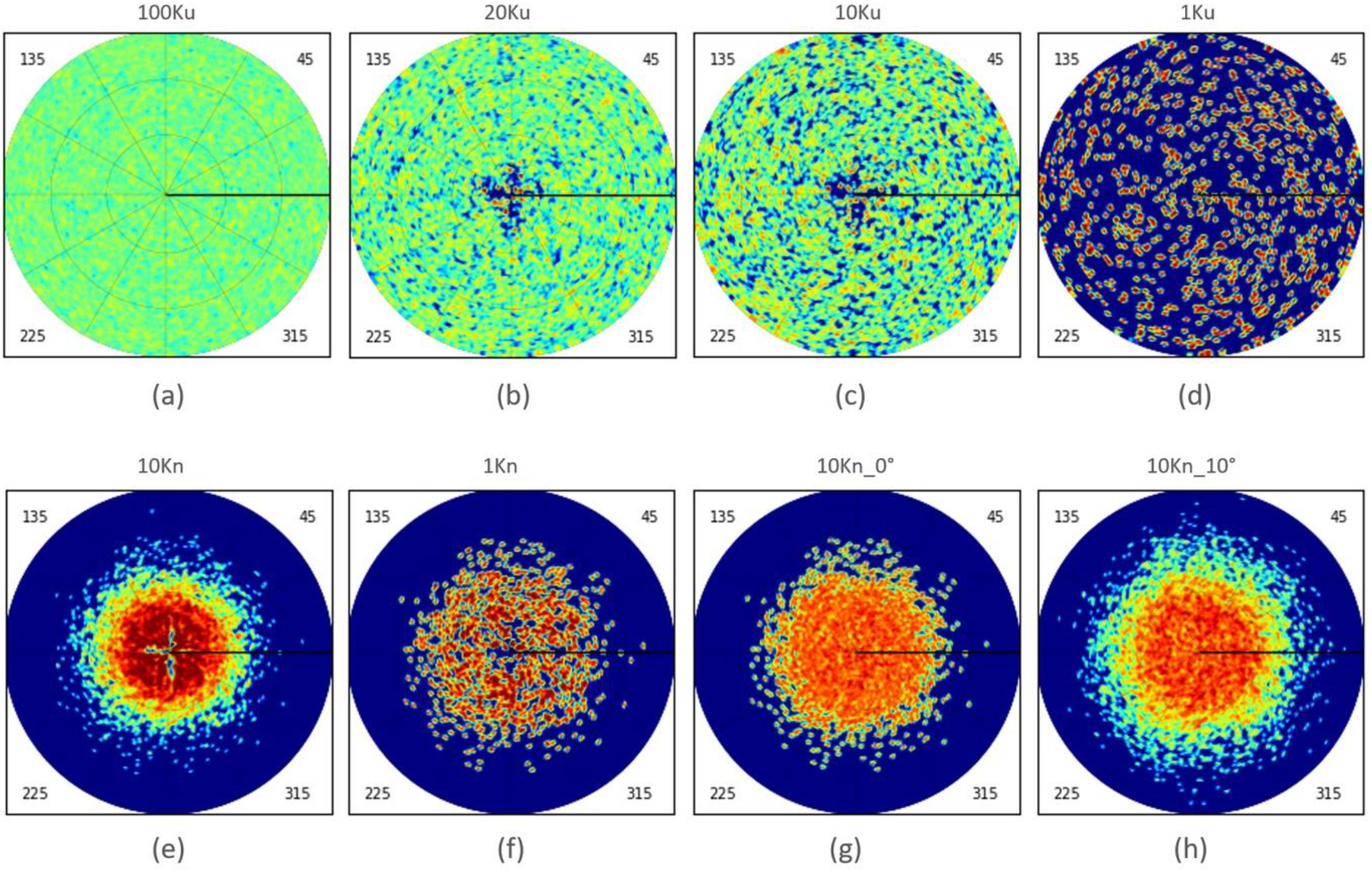
Distribution of views for the simulated data set. a) Initial uniformly distributed set 100Ku containing 100,000 projections; b) control uniformly chosen selection 20Ku, containing 20,000 projections; c, d) uniformly distributed subsets 10Ku and 1Ku, containing 10,000 and 1,000 projections, respectively; e) non-uniformly distributed subset 10Kn containing 10,000 projections; f) 1Kn-subset (1,000 projections) of 10Kn after removing the most overrepresented views; g) 10Kn subset corrected by replacing ∼3,600 overrepresented views by ‘dummy’ copies of the underrepresented ones; h) the same as g) but with perturbation of the orientation for ‘dummy’ copies with the mean 10°. Views distribution were plotted with *VUE*; the color scheme is the same as in Fig. 1.

Using the same atomic model, we calculated Fourier coefficients at a resolution up to 2 Å from the exact distribution of the electrostatic scattering potential (Peng, 1999) and computed a map using Fourier Transform (FT) with these coefficients (map noted as 2ÅFT).

### 2.5. Tools to analyze map quality

#### 2.5.1. FSC curves

To estimate quantitatively the quality of the 3D reconstructions, we used several measures including the commonly used FSC curves (Saxton & Baumeister, 1982; van Heel *et al*., 1982), fully understanding that its values (rather, uncertainty in its values) may be affected by the total number of projections. Such curves show consistency of the Fourier coefficients and therefore the confidence of the map details. At the same time, this does not exactly define the size of the details seen in the map, although it is relevant to it (*e*.*g*., Afonine *et al*., 2018, and references therein). For the tests with simulated data, we calculated the FSC between the reconstructed and the control maps as a measure of the closeness of the former to the latter.

In cryo-EM, the FSC curves are traditionally computed and plotted as a function of resolution expressed in Å^-1^. Calculations in the respective uniform scale lead to a strong statistical imbalance between the number of the Fourier coefficients per bin which varies drastically with the resolution. Figure 3a illustrates this for the 80S ribosomal data set described above. For an equilibrated binning, we introduced the scale uniform in Å^-3^ (Fokine & Urzhumtsev, 2002; Liebschner *et al*., 2019; Urzhumtsev, 2025b). Bins with the bounds uniformly chosen in such scale contain a roughly equal number of data (Fig. 3b), which is statistically more appropriate. Another advantage of this type of scale is that it zooms on the right-hand part of the curve, the region of high spatial frequencies, which is often the resolution interval of interest.

**Figure 3.**
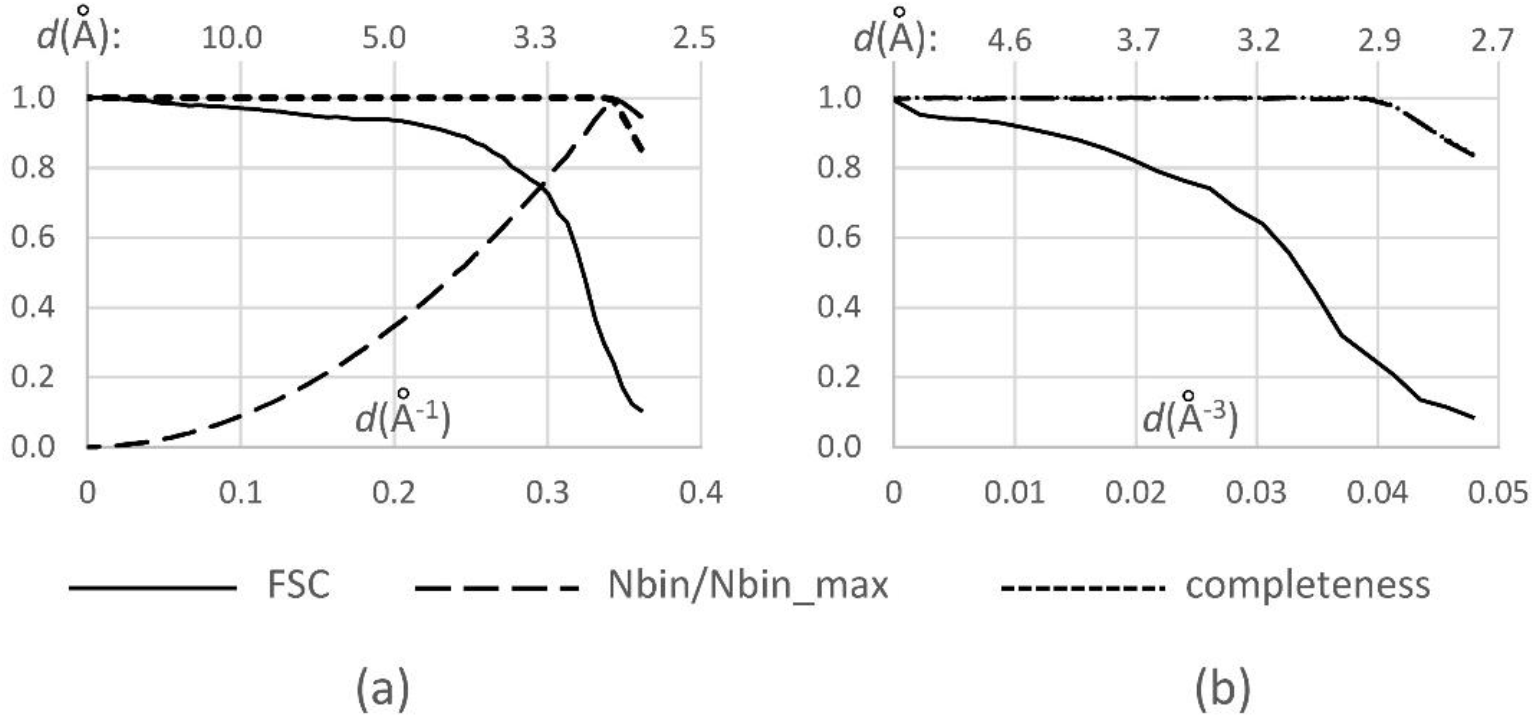
FSC curves for the 80S ribosome half-maps shown in different resolution scales. a) FSC curve calculated as a function of resolution in the conventional scale uniform in Å^-1^; the dashed line illustrates a tremendous variation of the number of Fourier coefficients per bin, *N*_bin_, shown as a ratio to their maximal number, *N*_bin_max_. b) the same curve calculated in the scale uniform in Å^-3^ giving a near equal number of the Fourier coefficients for all resolution shells with the full completeness (shown by the dotted line which, in b), coincides with the dashed line).

#### 2.5.2. Map discrepancy *D*-function

Whatever the resolution scale is, the FSC curve is a characteristic of the map Fourier coefficients and not of the map itself. To characterize how much two maps, calculated on the same grid, are close to each other, we used the map discrepancy function, a measure which corresponds to a visual map comparison (Lunin, 1988; Urzhumtsev *et al*., 2014).

First, one chooses the percentile value *p* = *N*_*selected*_/*N*_*grid*_ of the unit cell volume, where *N*_*grid*_ is the total number of the grid nodes in each of the two maps. Then, for the given couple of maps, we identify the cut-off levels *ρ*_1_ and *ρ*_2_ which select the respective number *N*_*selected*_ of the grid nodes with the map values above the respective cut-offs. Third, we analyze the difference between the selected regions by counting the number *N*_*diff*_ of grid points by which the masks of these regions differ. Finally, we normalize the computed value with respect to the theoretical number obtained for comparison with a random-valued map. By varying the percentile *p*, we obtain the discrepancy function *D* defined as

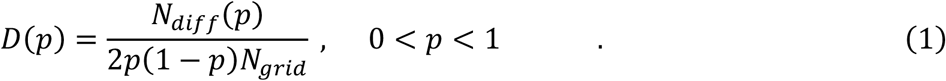

Values of *D*(*p*) close to zero mean a high map similarity at a given percentile *p*, while *D*(*p*) values close to one mean that the obtained maps are near random to each other considered at such particular percentile *p*. The regions of interest, those selected at 1σ – 2σ level in crystallographic maps, usually correspond to *p* values in the limits 0.01 – 0.05 (Urzhumtsev *et al*., 2014). Regions of interest about the macromolecule in cryo-EM maps are of order of magnitude smaller, in percentile of the unit cell volume, due to a much looser molecular packing in the virtual unit cells.

### 2.6. Control 3D reconstruction for simulated data

Before computing the control 3D reconstruction from the simulated 2D projections, we used the proposed validation tools to compare the two simulated maps, 2ÅC and 2ÅFT, defined in Section 2.4. As expected, they were different from each other (Fig. 4a). In particular, FSC between them decreased up to 0.7 when the resolution approached to 2 Å. A similar effect was observed for two other control maps calculated by *Chimera* and by FT at 5 Å and at 10 Å resolution.

**Figure 4.**
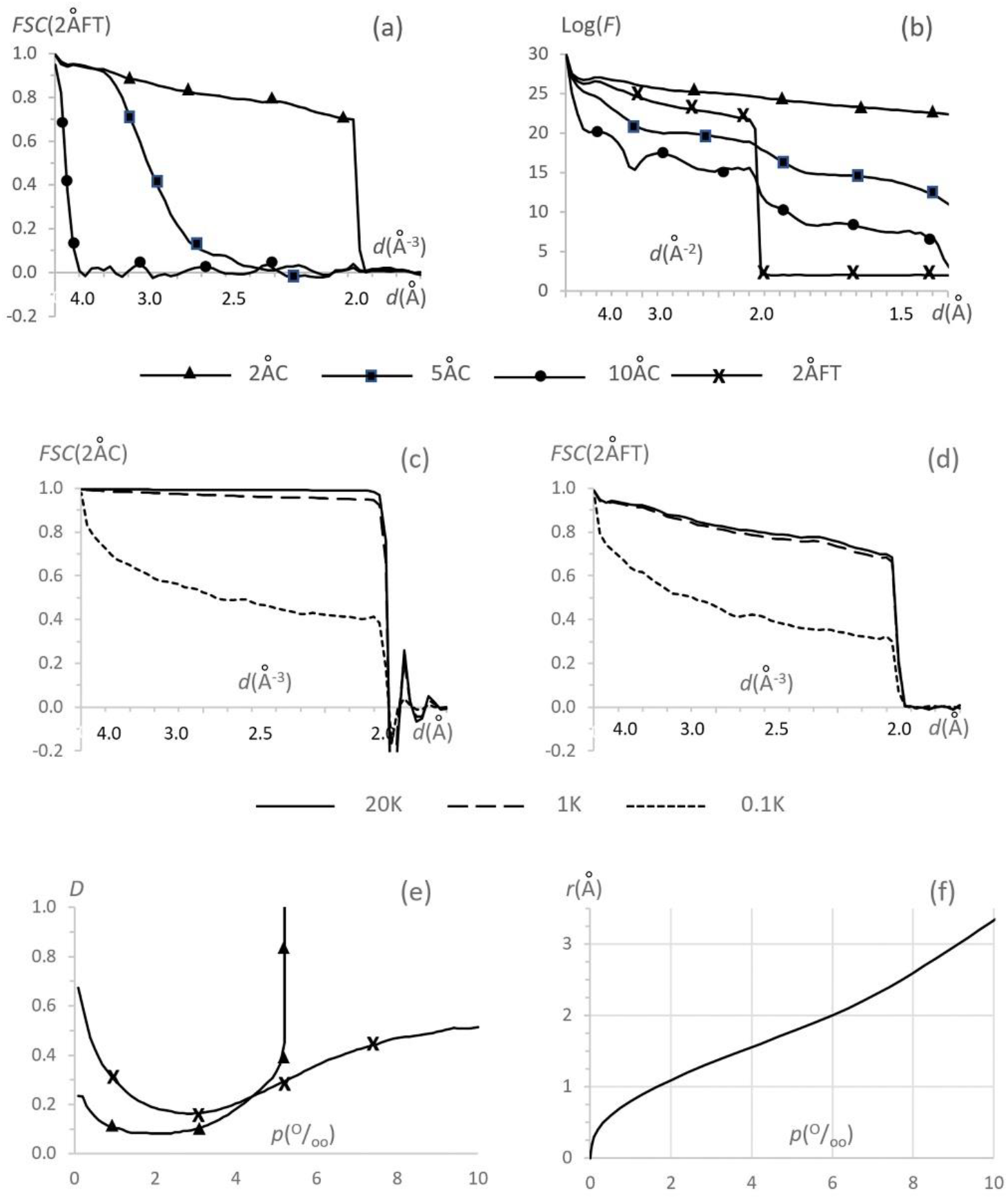
Analysis of the control maps. a) FSC curve between maps of different resolution calculated by *Chimera* and the 2 Å resolution map at calculated with FT; the resolution scale is uniform in Å^-3^; b) Logarithm of the mean intensity of the Fourier coefficients as a function of resolution calculated for the same maps as in a); the resolution scale is uniform in Å^-2^; c) FSC curve between the maps reconstructed with different number of projections selected randomly and uniformly, and the 2Å resolution map calculated by *Chimera*; the resolution scale is uniform in Å^-3^; d) a similar comparison with the 2Å resolution map calculated by FT; e) *D*-function between the 2Å resolution maps calculated by *Chimera* and by FT and the 20K control reconstruction; f) atomic radius giving the molecular mask of the required volume; volume scale is in per thousand for both e) and f).

To understand such difference, we computed the mean intensity of the Fourier coefficients of these maps as a function of the resolution. The map calculated as a Fourier transform showed an expected sharp drop of the intensity of the Fourier coefficients at 2 Å to the noise level (Fig. 4b). However, the Fourier coefficients for all three maps calculated by *Chimera* were significantly different from zero at a resolution higher than 2 Å for (Fig. 4b). This can be explained if these maps were actually calculated as sums of atomic contributions taken as a ‘peaky’ function approximating the central peak of atomic images at the respective resolution, for example, the Gaussians (*e*.*g*., Diamond, 1971). In other words, blurring atomic images when decreasing the resolution was modeled by increasing *B*-factor values. The increased slope of the respective curves when decreasing the nominal resolution (Fig. 4b) corroborates with this hypothesis. Such Gaussian-type modeling allows fast computing of an approximate map while ignores the actual shape of atomic images at a given resolution, including respective Fourier ripples, which may lead to certain map distortions (*e*.*g*., Urzhumtsev & Lunin, 2022).

Then we made several 3D reconstructions from the generated 2D projections using the software *Relion* and the 10ÅC map as the reference object. Expectedly, using all 100,000 error-free projections for a small object was excessive. Instead, we calculated reconstructions with 20,000 projections or smaller, selecting them randomly and uniformly from this 100Ku set (Figs. 1b – 1d). Fourier coefficients of such reconstructions, done with several thousand of projections, were very close to those for the 2ÅC map (not shown). When the number of projections used was smaller than approximately one thousand, the coefficients of the reconstructions started to have significant errors. Figures 4c and 4d show this for the set composed of 100 projections. As a result of this analysis, we decided to consider the map calculated with 20,000 uniformly selected projections (referred to as 20Ku in what follows) as the reference, the best reconstruction expected. The maps calculated from a smaller number of uniformly distributed 2D projections, varying from 10,000 (10Ku) to 1,000 (1Ku), were used to simulate imperfect reconstructions yet acceptable.

To further analyze the 20Ku map that represents the best 3D reconstruction, we computed how much it differs from each of the two maps, the initial 2ÅC map and the Fourier-calculated 2ÅFT. The *D*-function shows that the highest valued points of 20Ku, roughly for *p* < 0.003, are close to those in both of these maps while slightly closer to the 2ÅC map (Fig. 4e). The points corresponding to *p* > 0.005, are closer to those in the 2ÅFT map. For the intermediate values, all three maps are relatively close to each other. If one calculates a model mask as a set of spheres of a given radius *r*_*atom*_ and centered on atoms, these percentile values of 0.003 and 0.005 correspond roughly to *r*_*atom*_ = 1.36 Å and to *r*_*atom*_ = 1.78 Å (Fig. 4f). Below, we refer to the mask with *p* < 0.004 (*r*_*atom*_ = 1.6 Å) as the molecular region.

These results agree with the fact that 20Ku reproduces the peaks around atomic centers, (accurate in the 2ÅC map) while it ignores the Fourier ripples which are present in the maps of a limited resolution (2ÅFT map). The ripples of a given atom modify the map at some distance from the center, while the peak values themselves are perturbed by ripples from the neighboring atoms (Urzhumtsev *et al*., 2022). Both these corrections are not reflected in the 2ÅC map and therefore in the 2D projections used to reconstruct 20Ku.

A visual comparison of these maps at conventional cut-off levels selecting the molecule (varying about 20 σ in the given maps) showed practically no difference between the initial 2ÅC map and both 20Ku and 1Ku reconstructions (not shown).

## 3. Results

### 3.1. Reconstructions with simulated data

#### 3.1.1. Reconstructions with non-uniformly distributed views

To analyze the effect of a non-uniform distribution of views, we generated two subsets of views of the full set 100Ku. The views for the first subset, referred to as 10Ku, were selected uniformly. Reconstruction with 10Ku resulted in a map giving an FSC with the reference map 20Ku practically equal to one for all shells with the resolution lower than 2 Å, the resolution imposed for the initial map. The FSC values are actually quite high even for higher-resolution Fourier coefficients which are non-zero for these two maps (blue curves in Figs. 5a, 5b).

**Figure 5.**
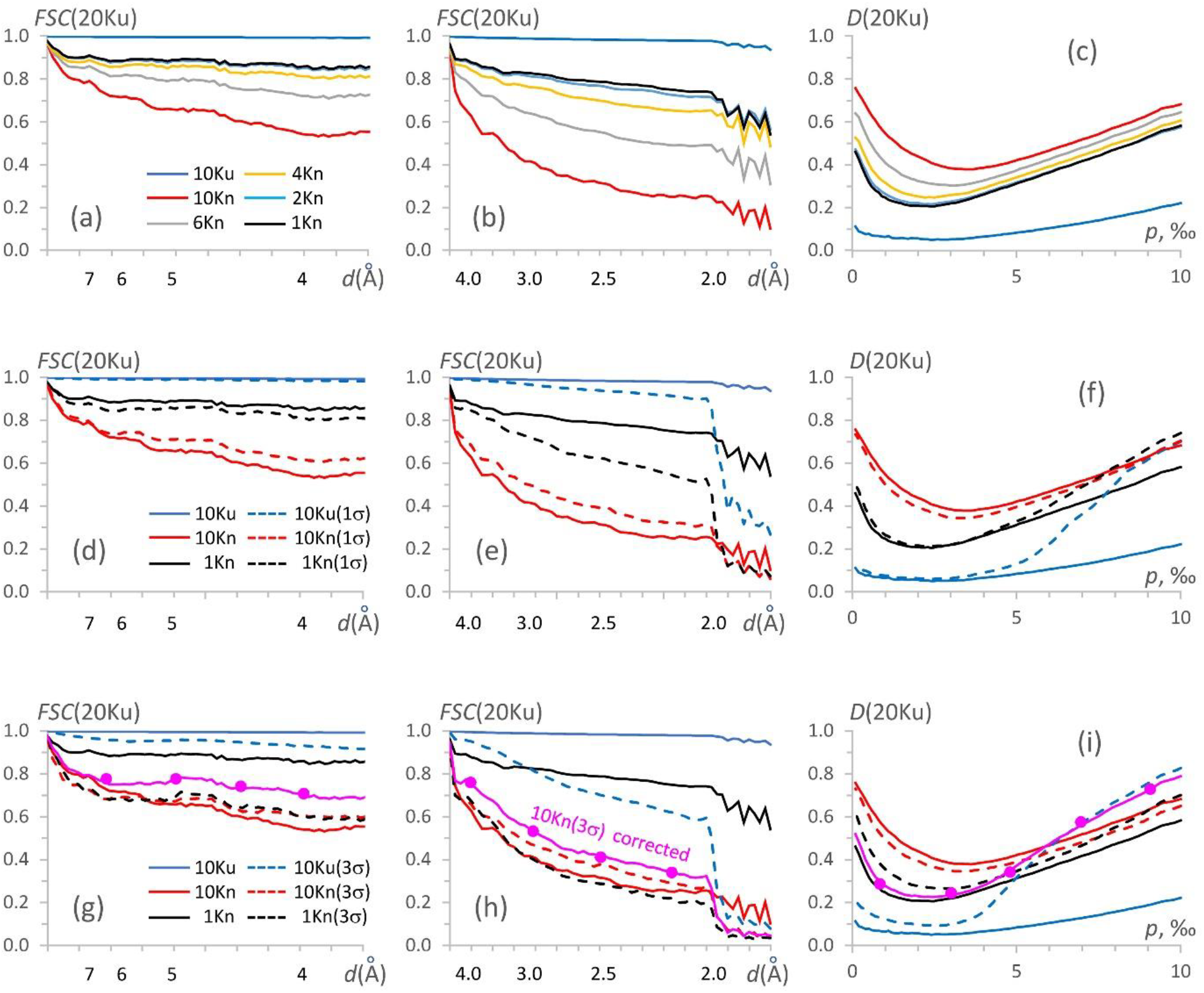
Data analysis for the IF2 simulated data. FSC curves calculated between different maps and the reference map 20Ku in the lower (left column) and higher (medium column) resolution intervals; the resolution scale is uniform in Å^-3^ while values in Å are given for reference. The right column shows respective *D*-functions as a function of the per thousand of the unit cell volume. Upper row shows the plots for the 10Ku map for the uniformly distributed 10,000 views (blue), 10Kn map for non-uniformly distributed 10,000 views (red) and the maps calculated with the reduced non-uniformly distributed sets of projections, up to 1,000 views (black). The medium row shows, by full lines, the results obtained with the error-free sets 10Kn and 1Kn, and, by dashed lines, the reconstructions with the same sets but with 1-rmsd errors in the projection values; the color code is the same as above. The bottom row is similar to the medium row but for the data with the 3-rmsd-error projection values. The curves in magenta, with markers, are for the set of projections with ∼5,000 severely overrepresented projections removed and being replaced by ∼1,000 ‘dummy’ projections for the most underrepresented ones.

The second subset, referred to as 10Kn, contained the same number of 2D projections but concentrated around one axis (*i*.*e*., a preferential distribution), chosen for simplicity as Oz. These views were selected from the 100Ku set according to the two-dimensional normal distribution. Figures 2c and 2e illustrate these subsets 10Ku and 10Kn. FSC between the 3D reconstruction with the 10Kn and the control 20Ku map drastically failed down, decreasing near-linearly in the inverse-cubic resolution scale (Å^-3^) both for higher- and lower-resolution shells (red curves in Figs. 5a, 5b). The map for the respective 3D-reconstruction appeared elongated in one direction (Fig. 6a), consistent with the existence of preferential particle orientations.

**Figure 6.**
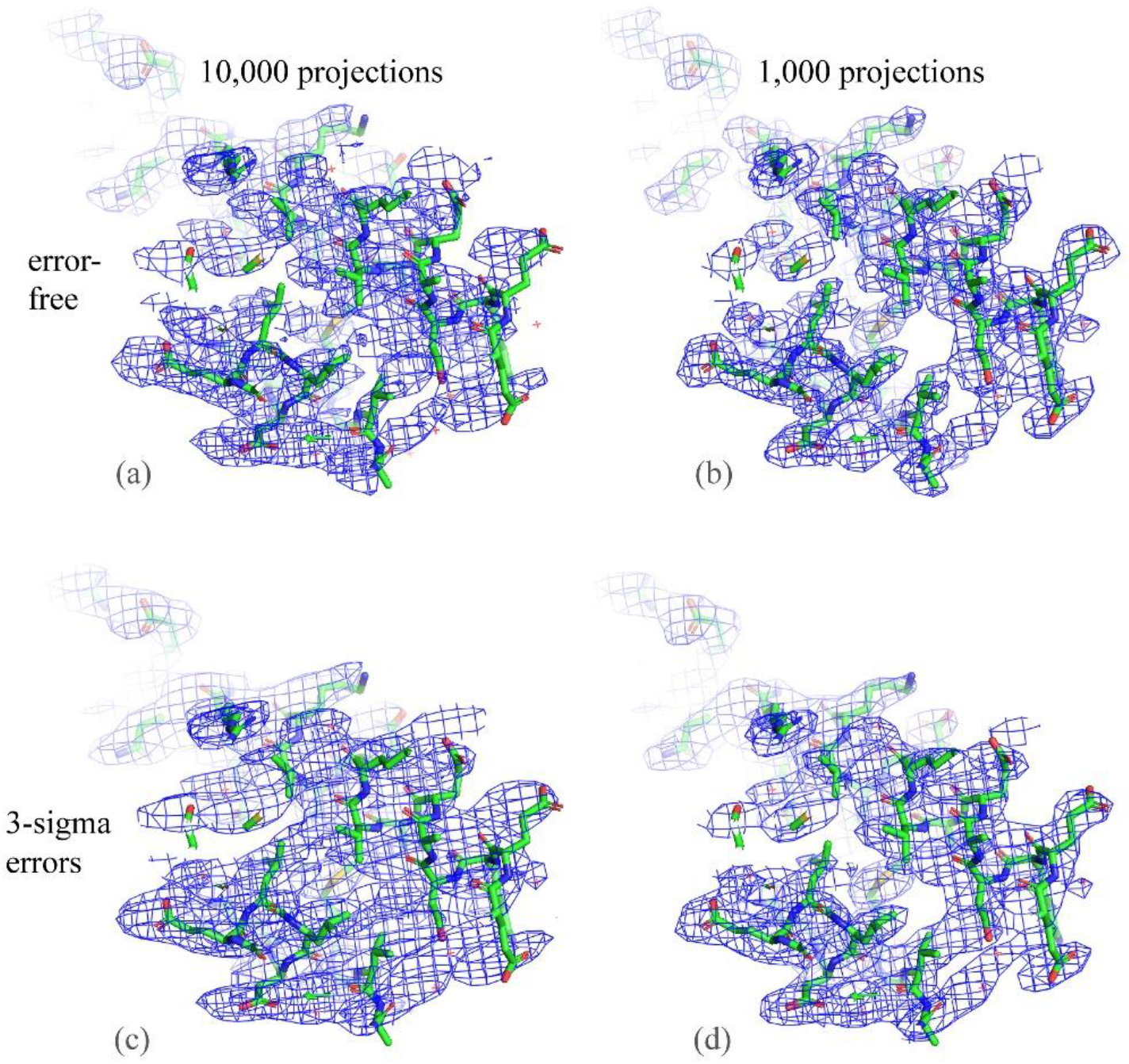
3D reconstructions for the IF2 simulated data. Reconstructions done with 10,000 non-uniformly distributed projections (left column) and with 1,000 (right column) after removing the overrepresented projections. Upper row shows the results of the reconstruction with the error-free 2D projections; the bottom row shows those with 3-rmds errors in the projection values. Maps in b) and d) are less stripy in comparison with those at a) and c), respectively, while d) shows some residual deformation in comparison with b). Figure prepared with *Pymol* (Schrödinger & DeLano, 2020).

The principal goal of the following tests was to improve, as much as possible, the latter 3D reconstruction by correcting the 10Kn set with no additional projections taken under new angles.

#### 3.1.2. Reducing overrepresented views, error-free projections

As a first approach, we equalized, as much as possible, the distribution of views in the 10Kn set by removing randomly the most overrepresented views, step by step, using the program *VUE*. Figures 5a and 5b illustrate the respective growth of the FSC with the reference map up to a certain limit; the improvement becomes marginal when reducing overrepresented views further from 2,000 to 1,000. For the subset with 1,000 views (1Kn) their distribution was near uniform except for empty regions (Fig. 2f). Overall, the FSC values comparing with the control 20Ku-reconstruction were quite high but lower than for the reconstruction with the 1,000 views distributed uniformly (1Ku; Fig. 5c). The reason for this is, in particular, a presence of empty angular regions, for which our procedure could not recover information, and we considered this result as the best possible under the given conditions. For these error-free simulated data, the deformation of the 3D reconstruction with the 1Kn subset of projections was rather marginal (Fig. 6b) especially for the principal regions of interest, corresponding to 0.002 < *p* < 0.004 (Fig. 5c), showing clearly molecular contours and main features.

This improvement of the results by equalizing the views distribution for error-free data sets by a simple reducing the number of overrepresented projections is not surprising. A large number of projections is important to filter over errors contained in 2D projections while this is irrelevant to error-free data. At the same time, information becomes lost when the number of projections falls below a certain limit.

#### 3.1.3. Non-uniformly distributed views with small noise

To analyze the effect of noise, we introduced artificial independent errors into the values of the 2D projections. They were introduced according to the normal law with the mean zero and sigma equal to 1 rmsd, the value calculated independently for each projection; in what follows, we refer to this set of projections as 1σ-projections.

First, we computed the 3D-reconstruction with the 10Ku data set of 1σ-projections, distributed uniformly, and compared it with the control reconstruction 20Ku. FSC values were close to those for the error-free data in the medium-resolution interval (Fig. 5d) and slightly lower in higher-resolution shells (Fig. 5e). The reconstruction inside the molecular region, corresponding to *p* < 0.004, was as good as for the error-free projections while significantly worse outside (blue curves in Fig. 5f).

For the subset 10Kn of non-uniformly distributed projections with noise, the quality of the 3D-reconstruction was similar to that for the error-free set (red curves in Figs. 5d, 5e). The reconstruction with the 1Kn subset of projections, after removing the most overrepresented views, significantly improved the FSC values. Nevertheless, the resulted FSC values were slightly lower than in the similar test with the error-free projections, especially for high-resolution shells (compare continuous and dashed black curves in Figures 5d and 5e with the respective red curves). At the same time, according to the *D* function, this did not affect the map accuracy inside the molecular region (Fig. 5f). Obviously, both maps were less accurate than the maps obtained with the uniform views distribution increasing *D* from approximately 0.05 - 0.10 to approximately 0.10 - 0.20 for *p* < 0.004 (compare black and blue curves). We conclude that for such relatively small errors in the projection values, 1,000 projections were sufficient to remove the noise by the respective averaging, and that, after removing severely overrepresented views, the residual map distortions were due only to the missed views.

#### 3.1.4. Non-uniformly distributed views with large noise

Then, we repeated the reconstruction but with the 3σ errors introduced into the projection values instead of the 1σ ones. For the uniformly distributed views, the FSC values for comparison with the control reconstruction 20Ku were close to those for the error-free data at a low and medium resolution but significantly worse at higher resolution, starting from approximately 4 Å (blue curves in Figs. 5g, h). Introducing strong but uncorrelated noise into values of the uniformly distributed projections, set 10Ku, does not distort much the reconstructed map inside the molecular region (corresponding to *p* < 0.004) but does it outside (blue curves in Fig. 5i).

Figure 6c shows the map reconstructed with 10,000 projections distributed non-uniformly with the 3σ-errors introduced into the projection values. For this reconstruction, the FSC values with the control 20Ku data were similar to those for the test with small errors when using the 10Kn sets (Fig. 5, red lines). However, in contrast to calculations with the error-free and 1σ-noise projections, leaving only 1,000 projections from 10Kn with large noise did not improve the FSC at medium resolution (Fig. 5g) and, just the opposite, made it worse at a resolution of approximately 3.5 Å and higher (compare black and red dashed curves in Fig. 5h). Interestingly, the *D*-function values slightly decrease, indicating some map improvement (Fig. 6d), even not as much as for similar calculations with 1σ errors (black and red dashed curves in Figs. 5f, 5i). We hypothesize that this reflects a slight improvement of the Fourier coefficients at medium and low resolution (Fig. 5g), that define the overall shape of the molecular image.

The plots in Figs. 5g, 5h show that the main errors in the recovered Fourier coefficients of a medium resolution come mostly from the non-uniform distribution and not from the noise in the data, while for higher resolution it is about half-on-half.

The results of calculations with error-free, 1σ- and 3σ-errors projections illustrate that in the latter case an insufficiently large number of projections leads to significant residual errors. This imperfection may be crucial when refining coordinates and *B*-factor values of atomic models versus cryo-EM maps. As a consequence, a simple removal of overrepresented views may by a not optimal protocol.

#### 3.1.5. Reconstructions completing underrepresented views

Trying to both equilibrate the views distribution and keep their number sufficiently large, we combined removing overrepresented projections with introducing some ‘dummy’ projections to complete underrepresented views, as described in the Methods section. Figure 7a shows the number of such views applying different values of the frequency cut-off level *ν*_*cut*_ (equilibrium frequency; Section 2.2). The ‘dummy’ projections (saying more correctly, extra references to the existing projections) were added with the exact values of the Euler angles describing the direction of the projections, or introducing random perturbations in these values, different in different tests, from small mean values up to a relatively huge value of 10° (Figs. 2g, 2h).

**Figure 7.**
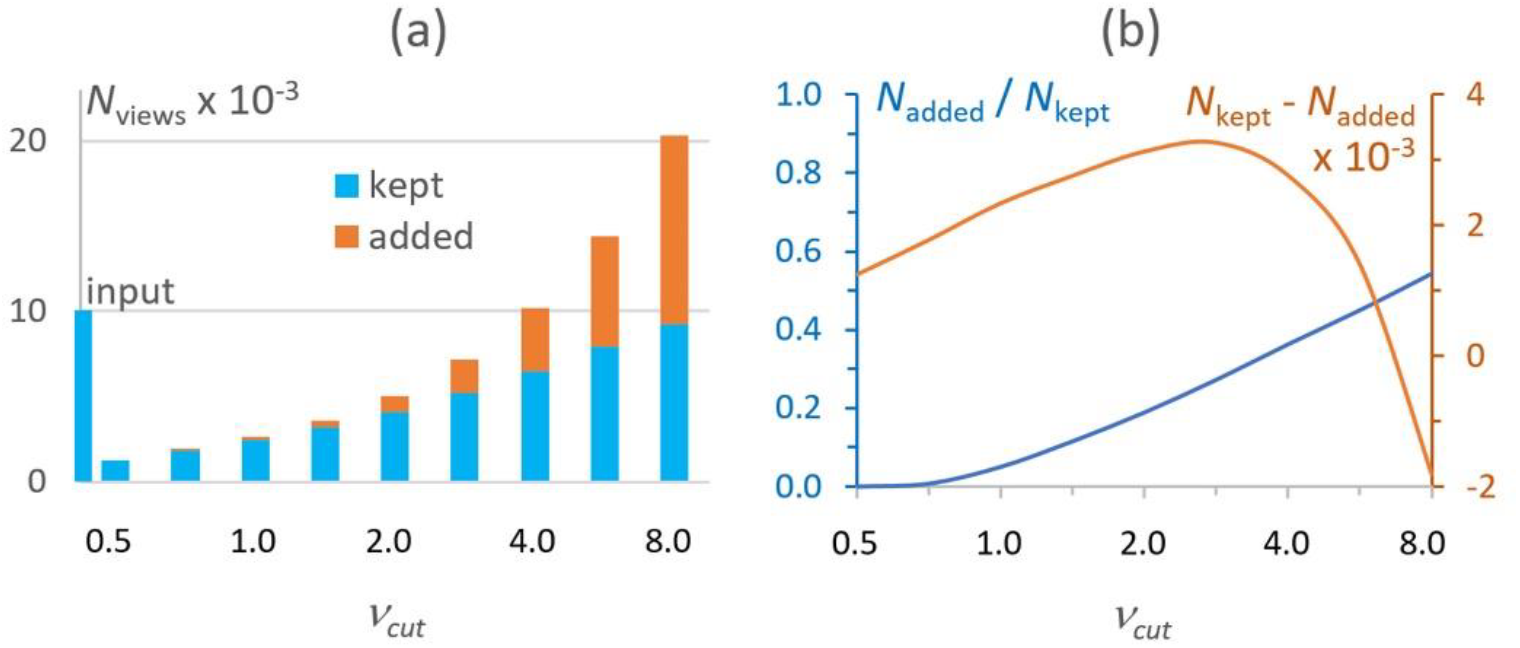
Number of views for the IF2 data as a function of the equilibrium frequency. a) Number of kept and added views, in thousand, to improve the non-uniform views distribution. b) Ratio and the difference of the added views with respect to the kept ones. Both plots are given as a function of the frequency cut-off *ν*_*cut*_ (uniform logarithmic scale).

The mostly improved 3D reconstructions have been obtained when the total number of views was either the same as before correction, *i*.*e*., about 10,000, or slightly reduced, up to about 5,000. For the given data set, this corresponded to the cut-off parameter values in the range 2 ≤ *ν*_*cut*_ ≤ 4, *i*.*e*., equalizing by the frequency 2 – 4 as calculated for the current distribution (Fig. 7a). This corresponds to higher values of the difference between the number of kept and added ‘dummy’ views (Fig. 7b). While the reconstruction with 10,000 total views gave slightly better FSC values (by 1-2%), the reconstruction with about 5-6 thousand total views gave the values of the *D*-function better by 2-3% (curves in magenta in Figs. 5g-i). Naturally, these estimates are only indicative, can be used for reference, and the optimal choice depends on a particular data set of projections and a particular structure. Figure 5i shows also that in the region of the molecule, *p* < 0.004, the reconstruction with such combined set of projections results in a map of similar quality as the best one that can be obtained using the error-free non-uniformly distributed set, namely, 1Kn set (curve in magenta compared with the black continuous curve).

Introducing artificial deviations in the values of the Euler angles for the generated ‘dummy’ projections, trying to better cover regions with the projections missed, did not improve the results but made them marginally worse. We can speculate that the refinement protocol recovered the original Euler angles rather precisely, even starting from very large orientation errors, thus giving the false impression that the distorted projections could better reconstitute the Fourier coefficients corresponding to the missed projections.

### 3.2. Correction of the experimental data sets

#### 3.2.1. 70S ribosome from *Staphylococcus aureus*

The initial 3D reconstruction for the 70S *S. aureus* ribosome data set was done with all cryo-EM projections, approximately 582,000 in total, in the presence of three clusters of overrepresented views (Fig. 8a). The map corresponding to the 3D reconstruction with this data set shows some anisotropy (Fig. 8d). Removing about a half of the most overrepresented views (about 285,000 projections left) blurred these three clusters of the projections (Fig. 8b). This made the respective map more homogeneous (compare Fig. 8b with Fig. 8a). Nevertheless, the map was yet distorted in some directions, as previously (Fig. 8e).

**Figure 8.**
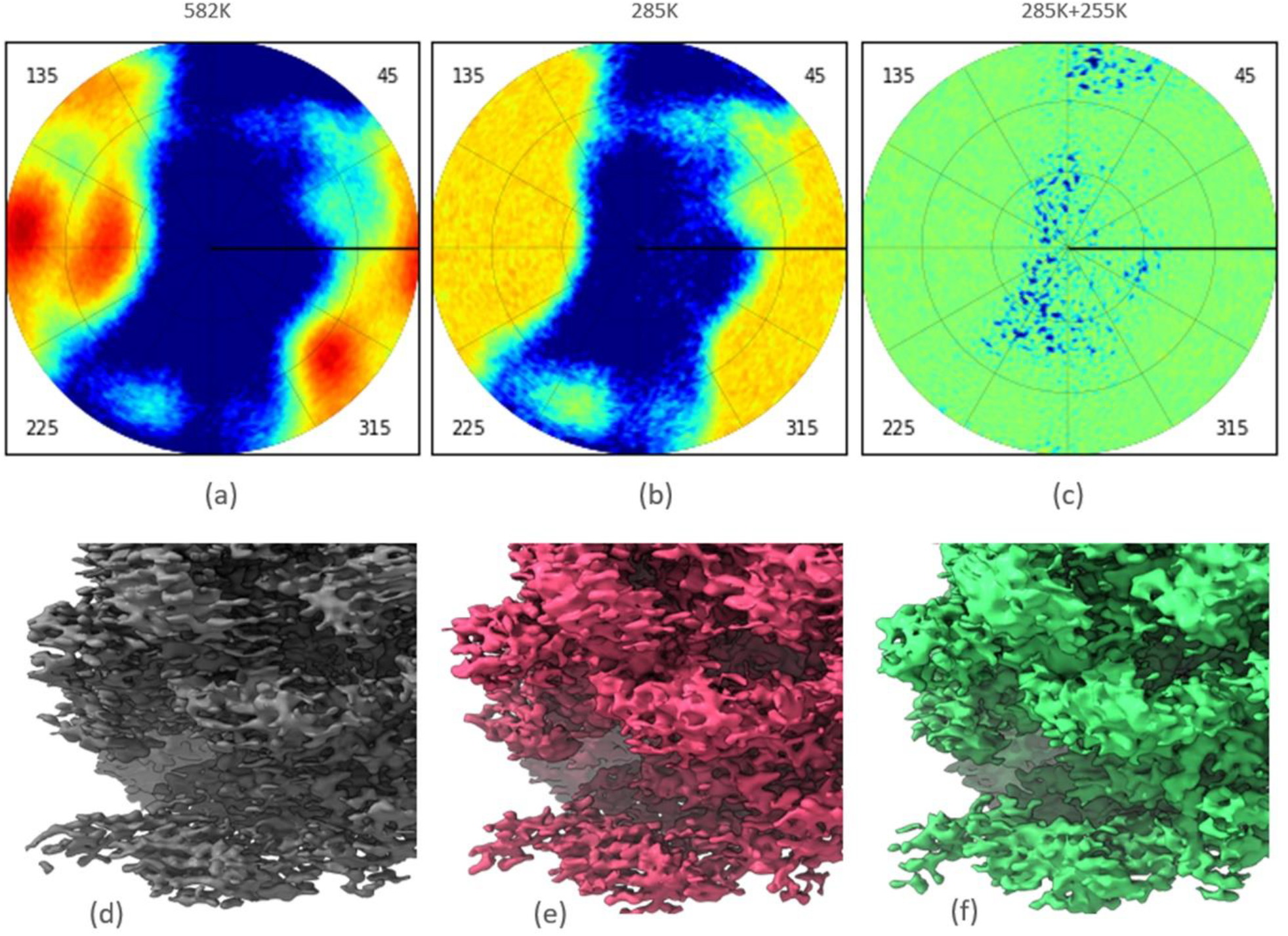
70S *S. aureus* ribosome data sets. Top row: views distribution plotted with *VUE*; a) Initial data set. b) Data set after removing a half of the most overrepresented views. c) Data set after removing a half of the most overrepresented views and completing by approximately same number of underrepresented views. The color scheme for the diagrams is the same as in Fig. 1. Bottom row: maps of the respective 3D-reconstructions. Maps were plotted with *Chimera*.

Finally, we removed the same number of overrepresented projections as in the previous calculation and completed the underrepresented projections by repetitive references to them, as explained in Methods, making the total number of views equal to ∼551,000. This time the angular space was covered nearly uniformly (Fig. 8c). This was possible because initially the set of views contained those for most of directions, while with very different frequency. Two opposite effects could be observed. On one hand, according to *Relion*, the nominal overall resolution decreased from 3.0 Å for the initial map to 3.1 Å for the intermediate map and to 3.2 Å for the last one. On the other hand, the final map was fully equilibrated, with no trace of the previous deformations or stripes in the images (Fig. 8f), and therefore easier to interpret.

#### 3.2.2. 80S human ribosome data

As shown in Section 2.3, a non-uniform distribution of views for the human 80S data set (Fig. 9a) leads to the 3D reconstruction being blurred anisotropically (Fig. 9e). Trying to improve the map, first, we reduced over-represented views removing randomly their extra copies. This left us with about 90,000 views. Their distribution was still significantly non-uniform (Fig. 9b) and the respective 3D reconstruction showed the same kind of defects as before (Fig. 9f).

**Figure 9.**
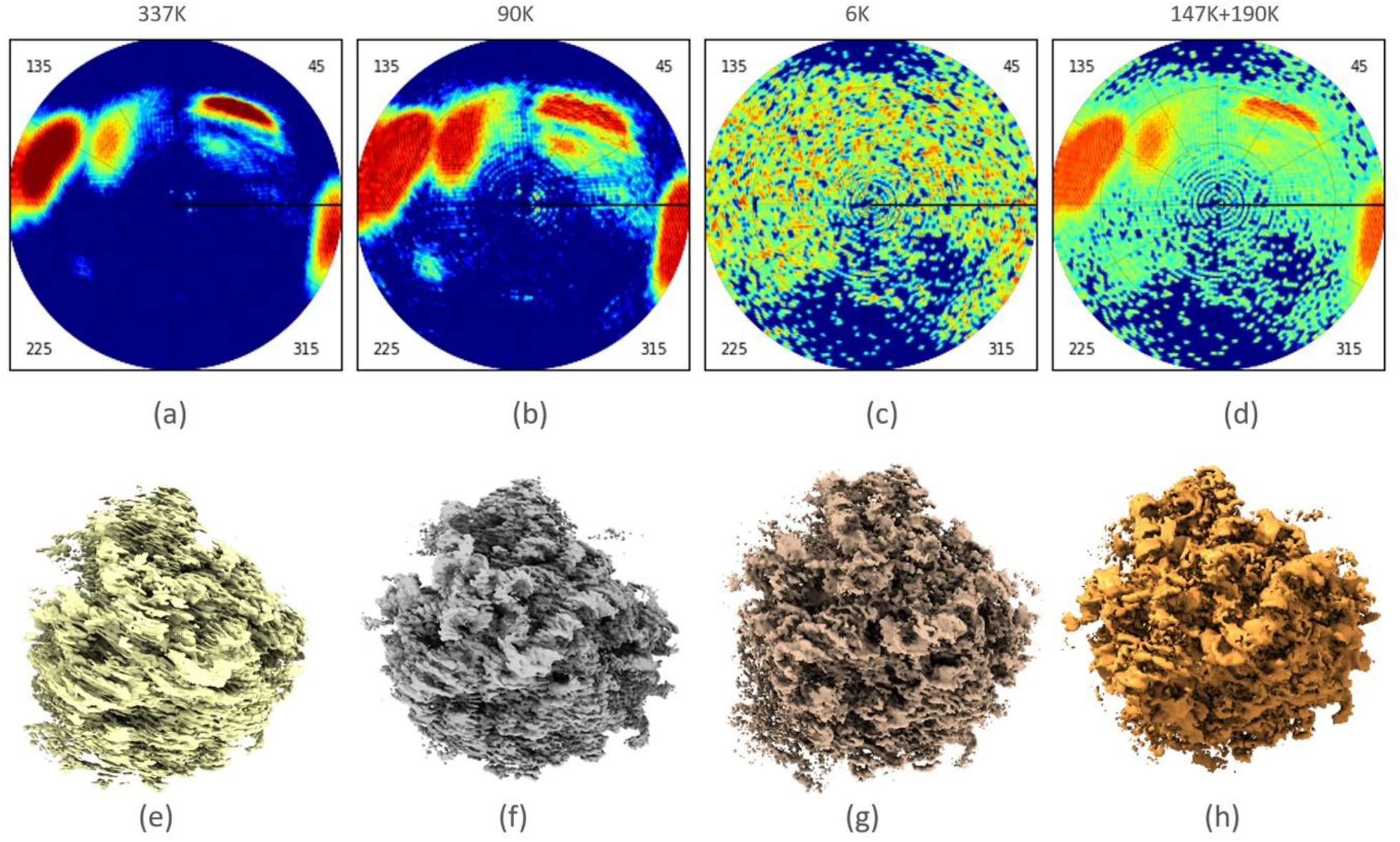
Human 80S ribosome data. Top row: views distribution plotted with *VUE*. a) Initial data set, about 337 thousand projections, the map shows characteristic stripes and distortions; b) its subset after removal of most overrepresented projections, about 90 thousand left; c,) the same, about 6 thousand left; d) the set composed of about 337 thousand projection after removal of about a half of the most overrepresented projections, and completing by a similar number of the underrepresented projections. The color scheme for the diagrams is the same as in Fig. 1. Bottom row: e) – h) respective maps plotted with *Chimera*.

Figure 9c shows the views distribution when we tried to equalize the distribution further, still by removing over-represented views even more. The corresponding set was composed of about 6,500 views only. While the corresponding 3D reconstruction was not blurred (Fig. 9g), the resolution was significantly reduced (*Relion* provided the value of 4.8 Å compared with 4.0 Å for the initial map).

Finally, we removed about half of the most overrepresented views and added a similar number of ‘dummy’ views corresponding to underrepresented projections taking each of them in several copies, according to their frequency. As discussed previously, this distribution was a kind of compromise between a higher number of experimental projections kept and an attempt to make the views distribution more uniform (Fig. 9d). This obtained 3D reconstruction was not deformed at all and contained more details (the *Relion* resolution estimate 4.0 Å) in comparison with the previous one and becomes much clearer to interpret (Fig. 9h).

## 4. Discussion

By their definition, irremovable errors both in crystallography and in cryo-EM cannot be corrected by any kind of refinement procedure and require a special treatment. For example, in crystallography, missing part of the atomic model can be modeled either theoretically (Lunin & Skovoroda, 1995; Bricogne & Irwin, 1996; Pannu & Read, 1996; Murshudov *et al*., 1997; Lunin *et al*., 2002) or explicitly by introducing dummy atoms (Isaacs & Agarwal, 1977; Lunin *et al*., 1985; Lamzin & Wilson, 1993). Some methodologically similar approaches may be suggested to correct irremovable errors in cryo-EM; severely non-uniformly distributed views are an example of such errors where an explicit introduction of ‘dummy views’ can be tried.

The results of 3D reconstruction depend both on the available set of the 2D projections and on the reconstruction procedure. We tested the most straightforward procedure using *Relion*, varying the number of references on a given set of the 2D projections, from zero, *i*.*e*. removing a reference to the respective projection, to several, *i*.*e*., artificially considering multiple copies of the same projection, with or without perturbating its view parameters, during reconstruction. Naturally, this can be seen as explicit weighting of such projections.

We observed that, as expected, for the projections with small errors in their values, a simple removal of the overrepresented views can help to some extent unless the number of the projections left becomes excessively small. On the contrary, for the data with significant errors, a simple removal of overrepresented projections is counterproductive. Removing the most overrepresented views completed by repeating information for underrepresented projections, thus making the final distribution almost uniformly distributed, results in 3D reconstructions with less deformations. Our tests suggest that such formal explicit correction of the set of projections should keep the total number of references close enough to that in the original set.

While map improvement after completing the underrepresented views by their extra copies is not always spectacular, new maps can facilitate model building when used alone or in parallel with the initial maps. Also, they may serve as a better reference map for further refinement of the 3D reconstructions, as well as for real-space refinement of atomic models.

To analyze the results of the reconstruction, we used the traditional measures like the FSC function with respect to the reference reconstruction. We observed that this information is not sufficient to reflect the actual similarity of maps and especially their appearance during visual analysis. For such goals, we used the discrepancy function *D* (Lunin, 1988), introduced earlier in crystallography.

Views distributions were analyzed with the program *VUE* (Urzhumtseva *et al*., 2024) which now also allows to routinely modify the lists of 2D-projections making the view distribution more uniform. Another advantage is that this program illustrates ‘on-the-fly’ these distributions. The program is available by request from one of the authors (AU) or from the Web site https://git.cbi.igbmc.fr/sacha/vue-cryo-em-software.

## Acknowledgments

We thank L. Fréchin for his help with test data. This work was supported by CNRS, Agence Nationale pour la Recherche (ANR), Fondation pour la Recherche Médicale (FRM), Institut National du Cancer (INCa), the Interdisciplinary Thematic Institute IMCBio, as part of the ITI 2021-2028 program of the University of Strasbourg, CNRS and Inserm, was supported by IdEx Unistra (ANR-10-IDEX-0002) and by SFRI-STRAT’US project (ANR 20-SFRI-0012), EUR IMCBio (ANR-17-EURE-0023) under the framework of the France 2030 program and LabexNetRNA (ANR-10-LABX-0036_NETRNA) administered by ANR. The electron microscope facility was supported by the Region Grand Est, FEDER, the French Infrastructure for Integrated Structural Biology (FRISBI) ANR-10-INBS-0005 /France 2030 program, EquipEx^+^ France-Cryo-EM (ANR-21-ESRE-0046) and Instruct-ERIC.

